# Putative linear motifs mediate the trafficking to apical and basolateral membranes

**DOI:** 10.1101/2020.07.13.200501

**Authors:** Laszlo Dobson, András Zeke, Levente Szekeres, Tamás Langó, Gábor Tusnády

**Author notes:** Contributed equally (Laszlo Dobson, András Zeke).

## Abstract

Cell polarity refers to the asymmetric organisation of cellular components in various cells. Epithelial cells are the best known examples of polarized cells, featuring apical and basolateral membrane domains. Despite huge efforts, the exact rules governing the protein distribution in such domains are still elusive. In this study we examined linear motifs accumulating in these parts and based on the results we prepared ‘Classical’ and Convolutional Neural Networks to classify human transmembrane proteins localizing into apical/basolateral membranes. Asymmetric expression of drug transporters results in vectorial drug transport, governing the pharmacokinetics of numerous substances, yet the data on how proteins are sorted in epithelial cells is very scattered. The provided dataset may offer help to experimentalists to characterize novel molecular targets to regulate transport processes more precisely.

## Introduction

Polarity is an essential feature of cells, especially in differentiated, multicellular organisms. In these cells, components (e.g. plasma membrane proteins, cytoskeletal components) are often organized asymmetrically. Many mammalian cell types exhibit a certain level of polarity, such as neurons, migratory cells, epithelial cells, and more. Epithelial cells possess a highly organized architecture establishing an apical-basolateral axis separated by tight junctions to maintain physiological barriers, as well as to deliver information to different regions of an organism [1], for example they maintain ion homeostasis in the eccrine glands and ducts [2] or play a role in nutrient uptake [3]. Although we have an increasingly growing knowledge of the main determinants of apical and basolateral polarity networks, the exact composition of these membranes are still elusive for most tissues [4]. Elements (proteins) required for the proper transport greatly differ on the apical and basolateral part of the membrane. In turn, polarity also relies on the correct sorting of these molecules to particular locations. Many times trafficking of these proteins from the Trans-Golgi Network to the plasma membrane does not occur in a single step, but rather via an indirect route through endosomal pathways [5]. During this journey to the cell surface proteins are tightly regulated via post-translational modifications and transient interactions with other molecules [6].

Many of these regulation processes are mediated via Short linear motifs (SLiMs), flexible protein segments composed of a restricted number of residues (usually between 3-10), that usually bind to ordered protein domains via coupled folding and binding. Their properties enable them to bind to a diverse range of partners with low micromolar affinity and establish transient interactions [7]. Besides mediating protein interactions, they also provide sites for post- translational notifications or proteolytic cleavage sites [8]. Recent decades provided a handful of evidence of motifs playing a crucial role in the trafficking of proteins to polarized membranes.

Trafficking to the basolateral and to the apical membranes include multiple pathways [9,10] and often include cargo sorting [11]. The basolateral targeting of transmembrane proteins may rely on cytosolic tyrosine [12], mono- and dileucine motifs [13,14]. Localization may also be proteolytic processing and glycosylation dependent [15]. In contrast, the apical targeting can occur in the absence of basolateral signal and may also involve rafts [16]. Both N- and O-glycans play important roles in apical sorting [17,18], as well as interaction between transmembrane regions and their surrounding [19]. Apical trafficking is sometimes functionally redundant or multipart, meaning proteins own a set of motifs, and those may serve as replacements for each other, individually capable of proper targeting [6]. The divided nature of apical membranes adds further complication to trafficking [20].

Although we have a moderate understanding which regions/residues/modifications play a critical role in individual proteins to reach their destination, the dozens of possible pathways makes it hard to apply general rules to them. Here we propose a novel approach to classify alpha-helical transmembrane proteins in polarized cells, based on their topology and putative SLiMs driving their localization. We collected hundreds of proteins with reliable experimental evidence of their destination and used computational biologist approaches to characterize motifs responsible for their trafficking. We used the resulting dataset to predict which membrane proteins localize to apical/basolateral membranes. Our dataset can be used by experimentalists to find new molecular targets governing transport processes in polarized cells.

## Results

### Ser/Thr rich motifs are abundant at the extracellular loops of apical transmembrane proteins

We scanned 200 human transmembrane (TM) proteins (100 apical and 100 basolateral) proteins for linear motifs, and then compared their distribution in disordered extracellular and cytoplasmic regions. To reduce the possible false positive hits, we only accepted putative motifs when conservation was visible across orthologs. We also random shuffled sequences ten times, and accepted motifs when the average plus three times the standard deviation was lower compared to real hits. We grouped each motif based on their localization, i.e. if they fall into a cytoplasmic/extracellular disordered region, or into any other segments. Next we counted the occurrence of motifs in apical and basolateral TM proteins (Figure 1).

**Figure 1.**
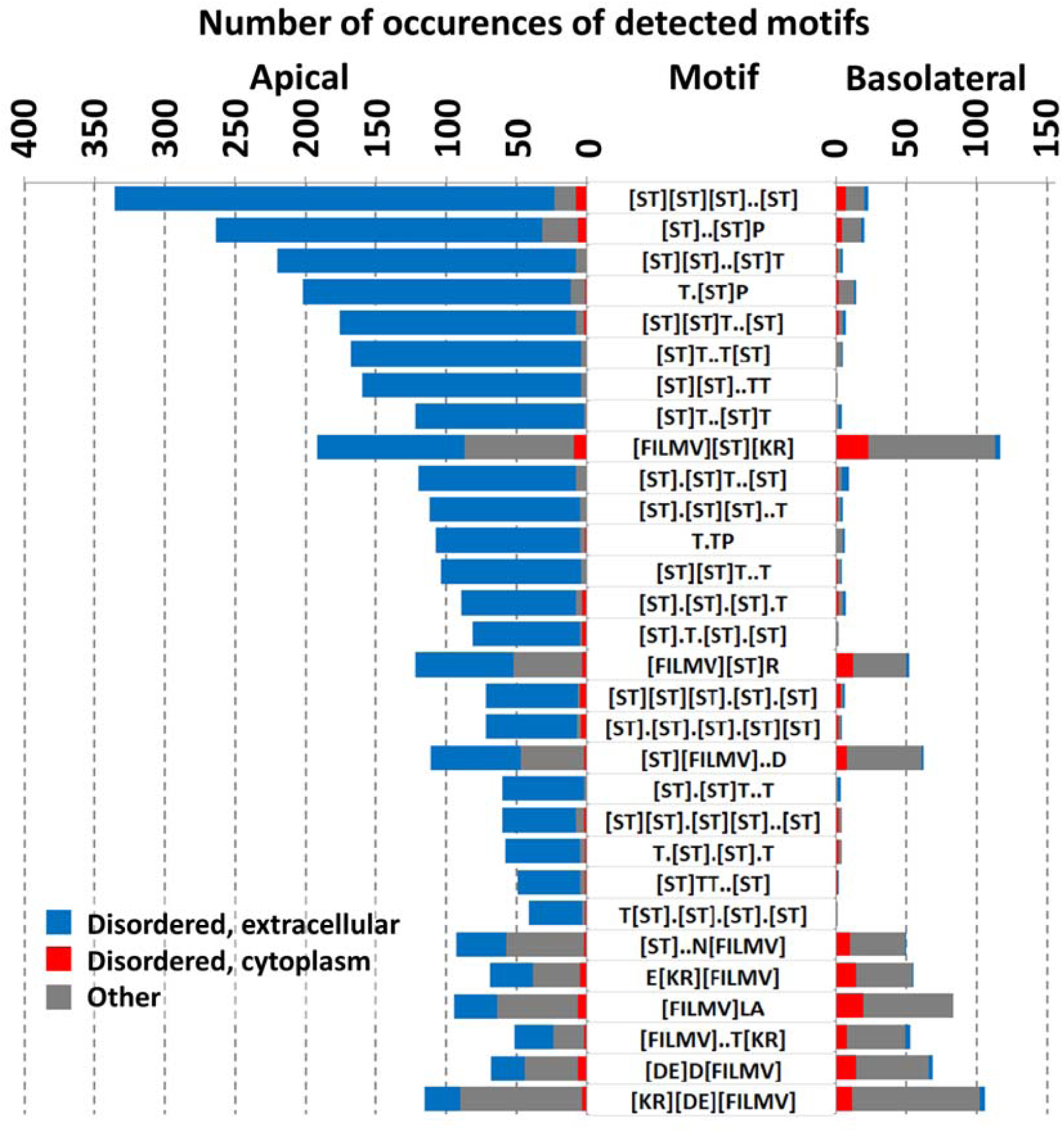
The first 30 most abundant putative linear motifs in apical and basolateral membrane proteins. We have to note, that these motifs highly overlap and they can be merged for a better consensus definition.

We found that motifs containing multiple Ser and Thr residues are highly abundant in the extracellular tail and loops of apical membrane proteins. Notably, ∼90% of these motifs are distant from the membrane region (>10 amino acids). Naturally, most of the identified motifs can also be merged for a better consensus description. At the second half of our list, hydrophobic and charged residue containing motifs also appear, however their discriminative power is relatively meager, compared to S/T motifs. The biochemical implications of the latter motifs are still unclear.

### Apical and basolateral membrane proteins can be classified based on the distribution of adjacent residue pairs

We prepared a ‘classical’ Neural Network (NN) to classify proteins based on their localization. Since apical/basolateral membranes can be considered as plasma membranes, we prepared four datasets containing apical, basolateral, plasma and other (Endoplasmic Reticulum, mitochondria, etc.) TM proteins. Input features included detected motifs and their extracellular/cytoplasmic localization, disordered and low complexity features, transmembrane topology and basic amino acid features. Although we achieved moderate success with this method (55% accuracy on multiclass classification), we concluded that there is still room for improvements.

We also prepared a Convolutional Neural Network (CNN), where protein sequences were converted into images. Each image contains 20×20 pixels, representing the 20 standard amino acids. Values in this matrix were calculated based on the distance of different residue pairs. Adjacent amino acid pairs increase the value of a point with a higher value compared to distant ones. The CNN achieved 61% accuracy as a multiclass classifier.

Last, we combined the output of the two predictors to classify proteins into 4 classes. By combining the output probabilities of the ‘classical’ and the convolutional NN, we increased the accuracy of the method to 66% (Note, as we have 4 classes, random prediction would achieve 25%). The most commonly occurring false cases were the cross-predictions of basolateral and plasma TM proteins, as apical and other TM proteins, respectively. Table 1 shows the binary evaluation of the method. In this case, there are 4 assessments for each group (apical vs other, basolateral vs other etc.). Using this evaluation the method performed quite well and achieved 78-85% accuracy.

**Table 1:** Binary evaluation of the proposed method.

|  | Training set |  |  |  | Validation set |  |  |  | Independent test set |  |  |  |
| --- | --- | --- | --- | --- | --- | --- | --- | --- | --- | --- | --- | --- |
|  | Apical | Basolateral | Plasma | Other | Apical | Basolateral | Plasma | Other | Apical | Basolateral | Plasma | Other |
| MCC | 0.56 | 0.72 | 0.65 | 0.78 | 0.56 | 0.68 | 0.6 | 0.75 | 0.51 | 0.47 | 0.38 | 0.58 |
| Sensitivity | 0.6 | 0.88 | 0.71 | 0.8 | 0.6 | 0.86 | 0.65 | 0.81 | 0.61 | 0.69 | 0.48 | 0.67 |
| Specificity | 0.94 | 0.87 | 0.93 | 0.95 | 0.93 | 0.85 | 0.93 | 0.94 | 0.89 | 0.81 | 0.88 | 0.9 |
| Accuracy | 0.88 | 0.87 | 0.88 | 0.91 | 0.86 | 0.86 | 0.86 | 0.9 | 0.85 | 0.78 | 0.78 | 0.84 |

Next we evaluated how the output probability (see methods) correlates with the accuracy (Figure 2). Predictions are sorted based on their probability value, the localization accuracies and reliability measured on the benchmark set are plotted against coverage. Reliability and accuracy correlates well. Half of the predictions have 70% or higher probability, where the prediction accuracy is 75%.

**Figure 2.**
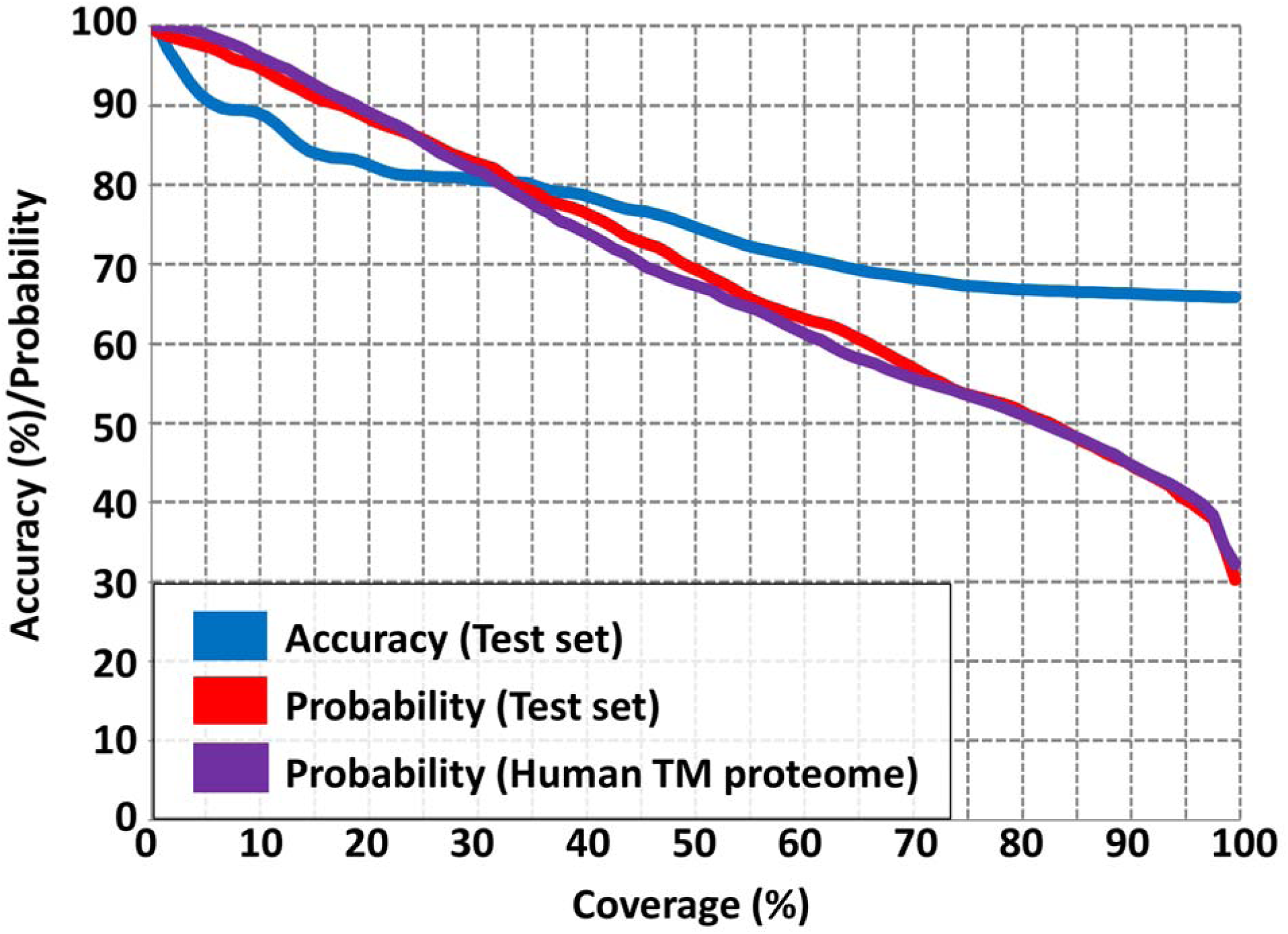
Correlation between the multiclass localization accuracy and probability. Predictions are sorted according to their probability values. Accuracies and the lowest probability measured on the subset of the benchmark set (blue and red, respectively) and probability values on the human transmembrane proteome (purple) are plotted against their rank in the sorted list. The Y- axis shows the coverage of the dataset (predictions above probability threshold divided by the number of the proteins in the benchmark set/TM proteome (coverage)).

### Dataset of apical and basolateral membrane proteins

We also run the prediction method on the human TM proteome. The classifier predicted 1285 proteins as apical, and 1475 proteins as basolateral. Considering the human TM proteome contains 5492 proteins, this means roughly 50% of these proteins may localize in the apical/basolateral membranes of epithelial cells during their lifetime. We have to note that cross-prediction from plasma/other to apical/basolateral TM proteins is more common than vice versa, thus the real proportion of such proteins is probably below 50%. We also measured the probability of the predictions on the full proteome and plotted it against accuracy and probability on the benchmark set (Figure 2). The probability distribution of the benchmark set and the proteome highly correlates, therefore it is safe to assume that the accuracy also shows a similar trend on the human TM proteome. Supplementary Table 1 contains the most reliable part of the prediction, i.e. predictions above 75% probability (381 apical and 649 basolateral membrane proteins).

## Discussion

### Examples with O-glycosylation regions governing sorting

While both apical and basolateral sorting can be complex and dependent on large inventory of different motifs, certain common features are already beginning to emerge. One of the most trivial features identified in our current study is the presence of O-glycosylation motifs or even complete regions in apically sorted proteins. For example, Sucrase-isomaltase (SI) is known to be sorted to the apical brush border in small intestine enterocytes, mostly governed by its O-glycosylated, membrane-proximal segment [21]. A similar, glycosylated “stalk” has also been found to govern the apical sorting of mucin1 in Madin-Darby Canine Kidney (MDCK) cells (representing the distal tubules of kidney) [22]. Such more-or-less extensive mucin-like regions also play a role in proper localization of the apical determinant podocalyxin, in addition to an NHERF-protein binding PDZ motif [23]. Although speculative, or results suggest that such regions might form the apical signal in many more proteins, such as maltase-glucoamylase. Accumulation of S/T rich motifs is likely connected with mucin-type O-glycosylation phenomena intimately connected to a preferentially apical sorting [21,23-24]. These sequences (especially due to their prominent Thr and Pro content) closely resemble the target sites of O-GalNac transferases [25-26]. However, due to the complex, processive nature of GalNac-T enzymes, the exact sequence of O-glycosylation sites is impossible to predict [27-28]. A cautious alignment of these regions reveal that although they are architecturally conserved within most vertebrate proteins, exact sequence matches are rare, as expected by the numerous imperfect, partially redundant O-glycosylation sites (Figure 3).

**Figure 3:**
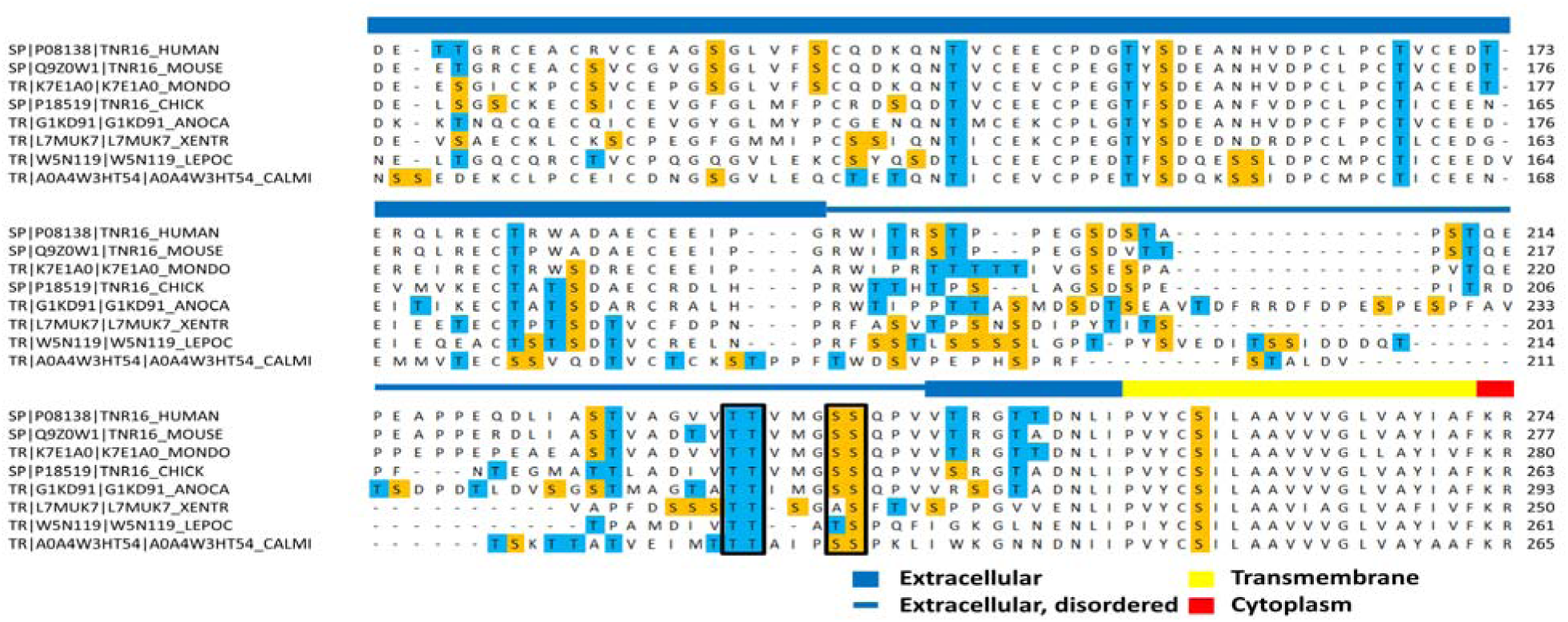
Sequence alignment of the O-glycosylated linker region between the extracellular domains (blue) and the transmembrane helix (yellow) of the low affinity Neurotrophin receptor (p75NTR or TNFR16). Despite excellent architectural homology between vertebrate receptors, only a few Ser-Thr rich repeats are consistently conserved. This figure was generated from an automated ClustalOmega aligment with slight corrections. HUMAN = Homo sapiens, MOUSE = Mus musculus, MONDO = Monodelphis domestica, CHICK = Gallus gallus, ANOCA = Anolis carolinensis, XENTR = Xenopus tropicalis, LEPOC = Lepisosteus oculatus, CALMI = Callorhincus milii.

Accuracy of disordered prediction methods is limited in TM proteins [29]. Although extracellular disordered regions are depleted in TM proteins [29], they are definitely present and one important function of them is to serve sites for glycosylation [30] mediating sorting.

### Limitation of our method

Despite the simplicity of our approach and its merits, this model also has many shortcomings. One of the most important problems relates to the fact that protein localization is not a binary variable in cells. What is more, apico-basal sorting of many proteins is not stationary, but depends on developmental stage of the cell as well as the actual tissue type. To circumvent these problems, our learning set was mostly based on proteins expressed or experimentally validated in mature, polarized MDCK cells, or tissues known to obey very similar sorting rules (e.g. small intestine enterocytes or the Caco-2 cell monolayers). However, there are other epithelial tissues whose sorting rules appear to be mildl (e.g. choroid plexus epithelium, with apical K+/Na+ pumps) or highly different (e.g. placental chorion epithel). Obviously, we need to learn much more of these specialized tissue polarities, before similar machine learning approaches become universally applicable.

Furthermore, current analytical methods provide limited information about polarized cells, as they usually characterize individual proteins with immunolocalization or with labeling amino acids selectively on one side of the polarized cell before proteomic analysis. All of these techniques provide a relative measure for the protein abundances in the different compartments of the polarized cell. The main bottleneck of immunolocalization is the limited availability of appropriate antibodies. For example, monoclonal antibodies can be highly specific and may only recognize one epitope, thus in this case we cannot safely assume that they are available in other regions of the polarized cell to the same degree. The most limiting factor in selective labeling that it uses primer amine specific reagents, enabling them to identify only those proteins that have such available regions. In accord, the coverage of TM proteins is relatively low in succeeding proteomic analyses, therefore differences between the two sides based on a limited number of peptides.

### Other resources, similar approaches

Recent decades provided several experimentally derived datasets that utilized high-throughput methods to classify localization of human proteins [31]. There are also a number of prediction methods that predict localization information, either the presence of a signal peptide [32], or the exact localization of proteins [33]. Some of these methods were trained on data automatically downloaded from computationally annotated databases, thus any bias in their sources affected their prediction accuracy, however, without any visible sign as their performance was measured on noisy datasets. In contrast we manually annotated each apical/basolateral membrane protein, this way we provided a very clean training set for our method.

Defining SLiMs is a rather laborious process, both by computationally and experimentally [34]. Dozens of pieces of evidence are required to confirm the functionality of a linear motif, as the information content of the peptide is very low and most of their occurrence in proteins are just ‘random’ false positive hits. There are several other efforts for the large scale computational identification and analysis of putative linear motifs in membrane proteins [35]. Here we demonstrated that such considerations have a strong predictive power, even if a significant proportion of the identified motifs is likely not functional.

The provided dataset can be directly utilized by experimentalists in the near future. We published a list of proteins that may have not been considered until now as potential targets and can be used to design selective inhibitors.

## Methods

### Training and testing data

We manually collected 171 apical and 125 basolateral membrane proteins. Although some of these proteins are available in high-throughput sets [31], each of these proteins was manually confirmed to be localized to the apical/basolateral membrane in kidney tissues. Furthermore, we collected 421 plasma membrane proteins from the RBCC database [36] and other sources [31], including our previous experimental pipelines [37–39]. We collected 521 proteins localizing to other membranes (Mitochondrial membrane, Endoplasmic reticulum, Lysosomal membrane etc) from UniProt [40].

We collected ortholog proteins from *Pan Troglodytes, Gorilla Gorilla, Pongo Abelii, Macaca Mulatta, Felis Catus, Canis Familiaris, Equus Caballus, Ovis Aries, Bos Taurus, Oryctolagus Cuniculus, Callithrix Jacchus, Mus Musculus, Rattus Norvegicus*, and *Sus Scrofa* from the OMA database [41]. We aligned the sequences with ClustalOmega [42]. We discarded those alignments, where we found discrepancies in the aligned transmembrane topologies.

### Prediction

We randomly selected 100 proteins from each localization group and used them for training and validating the predictions using 10/1 jackknife. Another 25 proteins from each group was used as an independent test set to measure the accuracy of the prediction. We used CCTOP [43] to predict transmembrane regions. We used IUPred [44] to detect disordered regions, however we masked out those segments that were also included in PFAM domains [45]. We used PlatoLoCo [46] to detect low complexity regions. Amino acid properties were derived from AAIndex [47]. Teiresias [48] was used to detect patterns in the sequence. We only accepted those occurrences, where the detected pattern fell into disordered non-membrane regions. We also checked orthologs and discarded non-conserved hits. We also random shuffled sequences ten times, and accepted motifs when the average plus three times the standard deviation was lower compared to real hits, similarly as described in the TOPDOM database [49]. Combinations of these features were used as an input to the classical NN. We used 16 input features, 8 hidden layers and stochastic gradient descent algorithm for the NN.

We also prepared a CNN, where each sequence was converted into a 20×20 matrix, and each point in the matrix represents a residue pair. The bottom triangle of the matrix represented extracellular disordered regions, while the top triangle represented cytoplasmic disordered regions. In each sequence, any neighbouring residue increased the point in the matrix by 2, while more distant pairs (up to 3 places) increased the point by 1. Each cell in the matrix was standardized using the same cells across the dataset. Ortholog proteins were also used during the training. The CNN consisted of 2 convolutional layers (filters: 32, dimensions: 3 and filters: 16, dimensions: 2) and a pooling layer (dimensions: 2) with relu activation functions and dropout (0.25). The final activation function was softmax with 4 output neurons.

To combine the results, first a vote was performed on the CNN prediction of different orthologs, using probabilities as weight. Finally, the normalized prediction accuracy of the classical NN and the CNN multiplied with their output probability was used as the final probability value.

### Dataset of apical and basolateral transmembrane proteins

We ran the prediction method described in previous sections on the Human Transmembrane Proteome database [50]. We selected the most reliable part of the prediction (above 75% probability, where prediction accuracy is 80%).

